# Genetic modulation of yield and phenotypic plasticity of yield in winter wheat

**DOI:** 10.1101/2024.11.08.622708

**Authors:** Nicolas Giordano, Victor O. Sadras, Mary J. Guttieri, Giovana Cruppe, Guihua Bai, Allan Fritz, Amy Bernardo, Paul St. Amand, Pablo Abbate, Andres Berger, Jeffrey Boehm, Amanda De Oliveira Silva, Sally Jones-Diamond, Scott Haley, Jane Lingenfelser, Guorong Zhang, Romulo Lollato

## Abstract

For winter wheat in the US Central Great Plains, phenotypic plasticity of yield is agronomically adaptive, *i.e.*, genotypes with higher plasticity have higher yield in high yielding environments with no tradeoff in stressful, low yielding environments. Using data from variety trials conducted between 2000 and 2022 and cultivars released between 1967 and 2022, we explored time trends in phenotypic plasticity and heritability of yield. We hypothesize that i) if yield plasticity is agronomically adaptive, then newer cultivars will have higher yield plasticity; ii) heritability of yield is declining in the time series; and iii) genomic regions associated with yield depend on the environment and do not fully overlap with those associated with phenotypic plasticity of yield. Breeding for yield and agronomic adaptation increased phenotypic plasticity of yield at 0.5% year ^-1^; broad sense heritability of yield decreased from 0.23 in 1993 to 0.15 in 2017. Genome-wide-association analysis shows genomic regions associated with yield varied between high yielding and stressful environments and were partially independent of those associated with yield plasticity. Newer cultivars have a higher frequency of alleles associated with yield and its plasticity. We discuss implications for breeding and agronomy aimed to improve wheat phenotypes.

## Introduction

Wheat (*Triticum aestivum L.*) grain yield is a complex trait that varies with genotype (G), environment (E), agronomic management (M) and their two and three-way interactions. Increased winter wheat yield in the U.S. Central Plains between 1985 and 2010 (Fischer et al., 2014) has been attributed to improved management and genetics and the interaction between of G and E whereby genetic gains have been higher in higher yielding environments (Cox et al., 1988; Berzonsky and Lafever, 1993; Donmez et al., 2001; Khalil et al., 2002; Fufa et al., 2005; Stewart et al., 2010; Green et al., 2012; Battenfield et al., 2013; Maeoka et al., 2020). Looking at the interaction between genotype and environment from the perspective of phenotypic plasticity provides links with ecology, evolution and developmental biology with implications for breeding and agronomy (Sultan, 1987; Sadras and Slafer, 2012; Sadras and Richards, 2014; Ruiz et al., 2019; Cooper and Messina, 2023; Giordano et al., 2024).

Phenotypic plasticity is “the amount by which the expression of individual characteristics of a genotype are changed by different environments” (Bradshaw, 1965); it depends on the trait, the genotype, and the environmental sources of variation (Bradshaw, 1965; Dewitt and Scheiner, 2004). Figure 1 illustrates genotype-dependent phenotypic plasticity of yield in a sample of wheat producing regions. In three out of six cases (Argentina, Australia, Alberta), yield plasticity was agronomically adaptive; this is, genotypes with higher plasticity had higher yield in high yielding environments with no tradeoff in more stressful, low yielding environments (Figure 1). A closer examination of the Canadian wheat belt shows plasticity was neutral for yield in Manitoba and positive in Alberta (Figure 1). In the Central Great Plains, one of the major wheat producing regions (USDA-NASS, 2024) and the focus of this study, previous investigations showed yield plasticity is agronomically adaptive (Grogan et al., 2016; Lollato et al., 2020a; Giordano et al., 2024). Thus, whether plasticity is agronomically adaptive depends on the environment, the germplasm pool (Figure 1), and the costs and limitations of phenotypic plasticity (DeWitt et al., 1998; Valladares et al., 2007; Auld et al., 2009).

**Figure 1.**
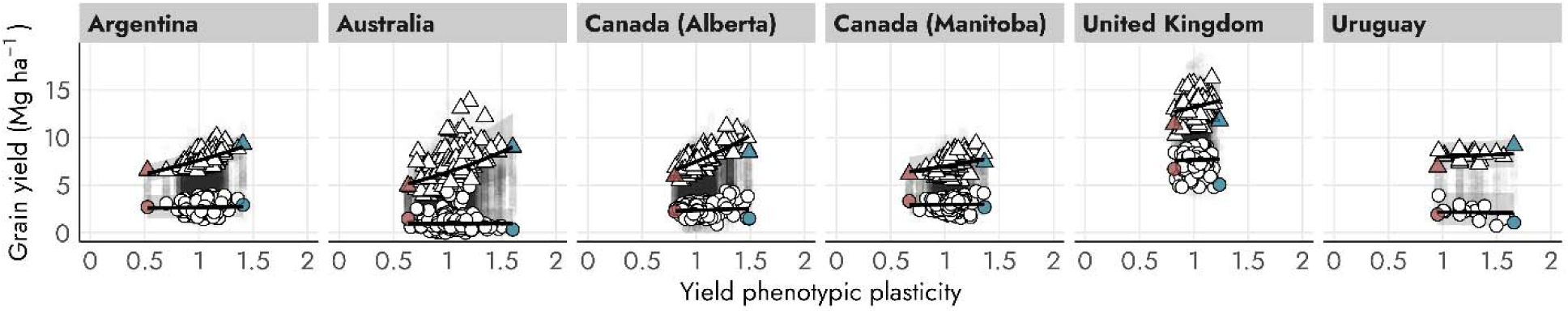
Association between genotype-dependent phenotypic plasticity of yield and actual yield in high (triangles, 95^th^ yield percentile) and low (circles, 5^th^ yield percentile) yielding environments in a sample of wheat producing regions. The genotypes with highest (blue) and lowest (red) plasticity are highlighted. Model used for quantile regression is in Equation 4. Data were obtained from variety testing networks in each region and reproduced with permission. Additional details on datasets used in this Figure can be found in Supplementary Table 1.

The rate of genetic gain in the yield of wheat is slowing down in major wheat producing regions (Brisson et al., 2010; Fischer et al., 2014; Lo Valvo et al., 2018; Maeoka et al., 2020). Common explanations for slower rates include diminishing returns and traits like harvest index approaching their biological limit (Hay, 1995; Fischer and Rebetzke, 2018). Declining heritability as a potential factor slowing down genetic gain has been overlooked; two perspectives, phenotypic plasticity and phenotypic connectance, converge to predict declining heritability of yield in some breeding settings. First, where selection for yield and agronomic adaptation favors higher phenotypic plasticity of yield (*e.g.*, Sadras and Lawson, 2011; de Felipe and Alvarez Prado, 2021), heritability might be declining because trait plasticity correlates negatively with heritability (Donovan et al., 2011; Sadras and Slafer, 2012; Alvarez Prado et al., 2014). Second, from the perspective of phenotypic connectance defined as the level of linkage between organs and traits in the developing organism (Amzallag, 2000), lower heritability has been predicted for a historic collection of Australian wheat for which phenotypic connectance of grain yield has increased over five decades (Sadras, 2024), and heritability and phenotypic connectance are negatively correlated (Amzallag, 2000).

Bradshaw (1965) formalized the notion that plasticity is a trait of its own, with its own genetic modulation and Scheiner (1993) advanced three models to explain the genetics of plasticity: i) the overdominance model, where plasticity is negatively related to the degree of heterozygosity; ii) the pleiotropic model, where genes have pleiotropic effects, influencing multiple traits simultaneously and iii) the epistatic model, which assumes the genetic independence of the trait *per se* and its plasticity. At present, the overdominance model is largely irrelevant in wheat varieties that are highly inbred and predominantly homozygous. Empirical evidence supports pleiotropic and epistatic models accounting for the partial genetic independence of phenotypic plasticity and the trait *per se*, *e.g.* for grain weight (Alvarez Prado et al., 2014) and grain yield in *Zea mays* (Kusmec et al., 2018) and for flowering time and fruit weight in *Solanum lycopersicum* (Diouf et al., 2020).

Using yield data of hard winter wheat collected between 2000 and 2022 from cultivars released between 1967 and 2022 in the U.S. Central Plains, we explore time trends in phenotypic plasticity and heritability of yield. Our working hypotheses are: i) if yield plasticity is agronomically adaptive, then newer cultivars will have higher yield plasticity; ii) heritability of yield is declining in the time series, and iii) genomic regions associated with yield depend on the environment and do not fully overlap to those associated with yield phenotypic plasticity.

## Method

### Field experiments

Grain yield data were obtained from hard winter wheat (HWW) variety trials conducted in the United States Central Great Plains between 2000 and 2022. During the last 10 years, the states of Kansas, Colorado and Oklahoma have sown about 6.1 Mha of HWW and produced an average 15.0 Mt (USDA-NASS, 2024) annually, with average yields from 1.8 to 3.7 Mg ha^-1^. This study analyzed the dataset comprising of 111,026 observations of 888 genotypes, including commercial cultivars and advanced experimental lines within breeding pipelines, hereafter both referred as cultivars, across 853 environments resulting from the combination of site and year; not all wheat cultivars were tested in all environments. The trials spanned over the states of Colorado, Kansas and Oklahoma (Figure 2A). Cultivars were released from breeding programs between 1967 and 2022. Details of field experiments can be found in Figure 2B and in Munaro et al. (2020).

**Figure 2.**
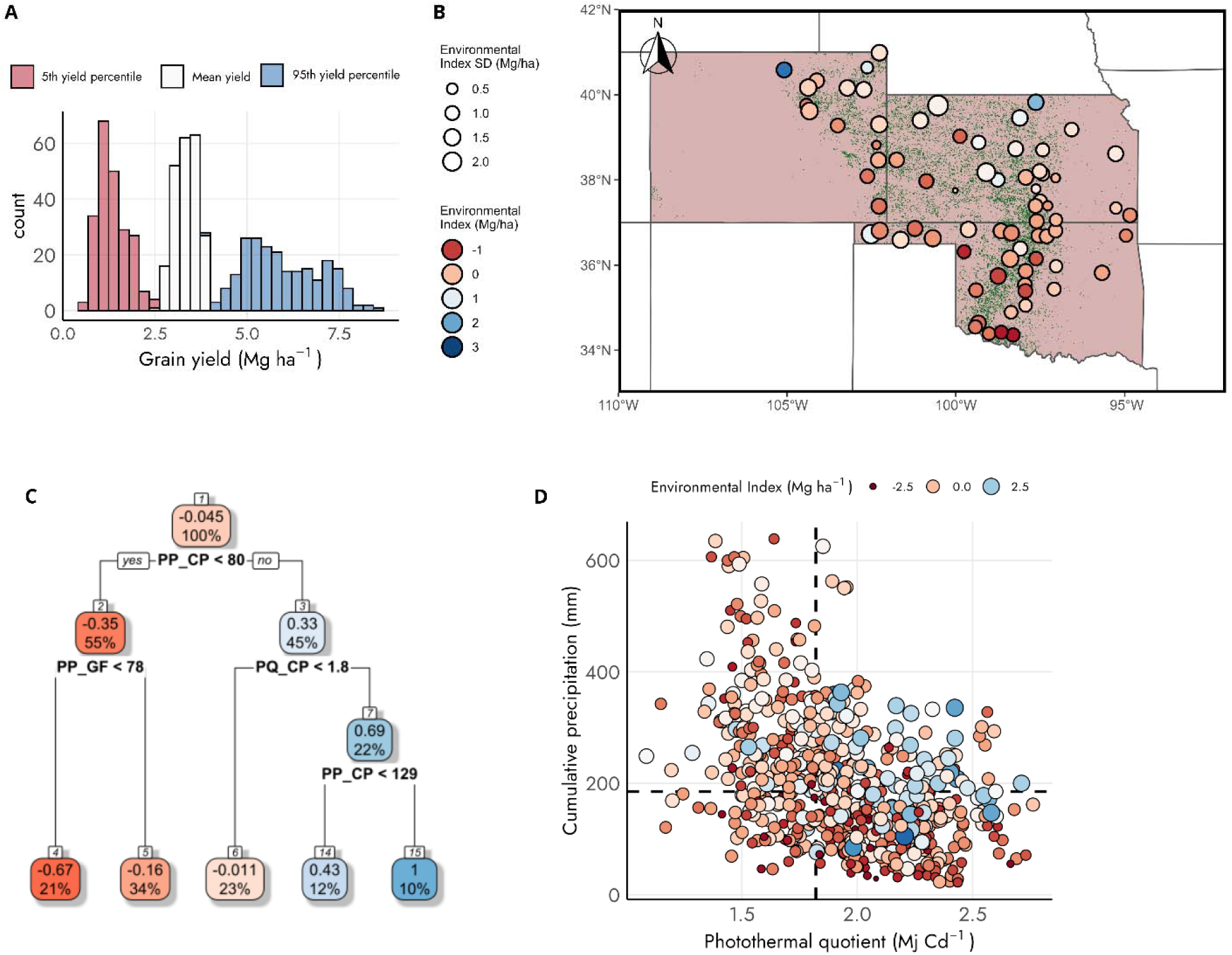
(A) Frequency distribution of grain yield in three levels: high yielding environments (blue, 95^th^ percentile), mean yield (white) and low yielding environments (red, 5^th^ percentile). (B) Spatial distribution of experimental sites throughout the states of Colorado, Kansas, and Oklahoma. In (B), each point represents an experimental site, color of the point is relative to the average environmental index of a given location, and their size represents the standard deviation of environmental indexes within a site. (C, D) Association between environmental index, , photothermal quotient and the cumulative precipitation from the start of the critical period (−300 °Cd from anthesis) and the end of the grain filling period (+600 °Cd from anthesis). Color of nodes/points is relative to the mean environmental yield. In B, C and D, environmental indexes are expressed as deviations from the overall intercept, (mean: 3.4 Mg ha^-1^, c.i.: 3.3 – 3.5 Mg ha^-1^), both estimated following Equations 1 and 2. Estimation of levels of the grain yield distribution shown in A are described in Equations 1,2 and 4. Abbreviations: precipitation (PP), photothermal quotient (PQ), critical period (CP), grain filling period (GF).

### Yield data

We filtered yield data to all cultivars replicated in at least 25 environments, since the predicted standard deviation of plasticity based on the exponential decay model was 0.03 with this number of environments (Supplementary Figure 1). This resulted in a set of 234 cultivars (Supplementary Table 2). We estimated yield phenotypic plasticity as the slope of a reaction norm (Woltereck, 1909) between observed grain yield and environment mean yield. This approach requires assumptions on the shape of the reaction norm but is less sensitive to uneven sample size from each genotype when compared to estimates from variance ratios that do not need these assumptions (Dingemanse et al., 2010). Yield plasticity was estimated with a Bayesian hierarchical approach (Equation 1, 2) using a Gibbs sampler developed for this purpose (Su et al., 2006) and implemented using the *FW* package available in R (Lian and Campos, 2015; R Core Team, 2021).

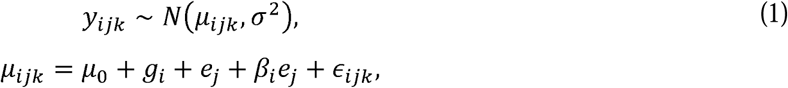

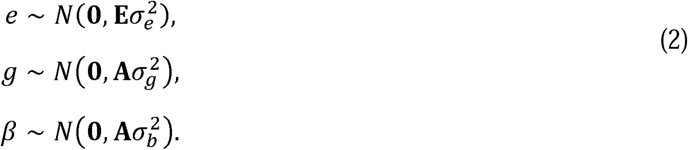

In Equation 1, *y_ijk_* is the observed yield at the *k^th^* replicate (*k*: 1,2, …, *n*), of the *i^th^* genotype (*i*: 1,2, …234) at the *j^th^* environment (*j*: 1,2, …234). The number of *n* replicates was on average four and varied with environment. In the process model *µ*_0_ is the overall intercept, while *g_i_* and *e_j_* are the main effects of genotypes and environments respectively, quantified as deviations from *µ*_0_. Thus, *g_i_* can be also interpreted as the intercept of the reaction norm and 1+ *β_i_* is the slope used to estimate phenotypic plasticity of yield. *∊_ijk_* is the error term assumed normally distributed with expected value equal to zero and variance 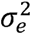. Prior distributions are all multivariate normal, **A** and **E** describe the covariance structure between levels of random effects *g_i_*, *β_i_* and *e_j_*, respectively (Equation 2). By retrieving variance components from model in Equations 1 and 2, we calculated broad sense heritability (*H*^2^) of grain yield as:

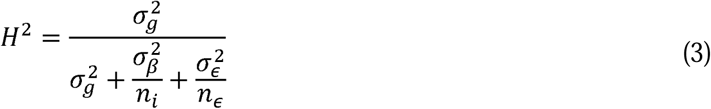

In Equation 3, 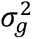 is the genotypic variance of genotype main effects (*g_i_*). The denominator in the equation is the total phenotypic variance, where 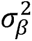 is the variance from the slope of the reaction norm (*β_i_*) and the residual error variance 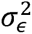, both scaled by their respective degrees of freedom *n_i_* and *n_∊_*. *H*^2^ of yield in the population was estimated using a sliding window of 10 years over the cultivars year of release starting in 1989 until 2022. Data before 1989 were excluded as only three cultivars were representative of the entire population in that period.

We explored how phenotypic plasticity of yield changed with cultivar year of release. A linear Gaussian model was used to assess the association of yield plasticity and year of release:

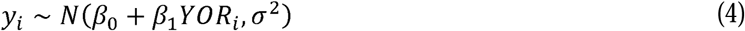

Quantile regression was used to estimate genotype-specific grain yield at high (95^th^ percentile) and low yielding environments (5^th^ percentile, Equation 5). The model was fitted using an asymmetric Laplace distribution, which allows specifying *τ* parameter to 0.95 for 95^th^ percentile and 0.05 for 5^th^ percentile:

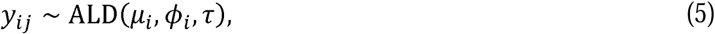

Where *y_ij_* is the observed grain yield at the at the *j^th^* replicate (*j*: 1,2, …, *n*), of the *i^th^* genotype (*i*: 1,2, …234), with expected value equal to *µ_i_* and the dispersion *ϕ_i_*.

Four yield phenotypes were then defined as: i) yield plasticity; ii) yield in low yielding environments (5^th^ yield percentile, *i.e., µ_i_* when *τ*= 0.05 in Equation 5); iii) mean cultivar yield (*g_i_*, Equation 2) and; iv) yield in high yielding environments (95^th^ yield percentile, *i.e., µ_i_* when *τ*= 0.95 in Equation 5).

The association between yield phenotypic plasticity and yield in low yielding environments (5^th^ yield percentile), yield mean and yield in high yielding environment (95^th^ yield percentile) were explored using linear regression:

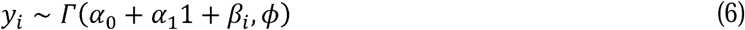

Where *y_i_* follows a Gamma distribution and represents the estimated value for each yield level for the *i^th^* genotype (*i*: 1,2, …234). The expected value is represented with *α*_0_ being the intercept and *α*_l_ the slope of the association of yield plasticity, *β_i_*, and three levels of yield, while *ϕ* is the dispersion.

### Environmental data

Daily weather data were retrieved from Daymet (Thornton et al., 2014) using the *daymetr* package available in R (Hufkens et al., 2018). Daymet provides daily weather gridded estimates of 1 km × 1 km spatial resolution from ground-based meteorological stations (https://daymet.ornl.gov/overview). Weather data were summarized for (i) the critical period for yield determination between −300°Cd and 100°Cd centered on anthesis using a base temperature of 4.5°C (Fischer, 1985; Cossani and Sadras, 2021), (ii) grain fill, from 100 °Cd to 600 °Cd after anthesis using a base temperature of 0°C (Dupont et al., 2006). Anthesis date was simulated from a locally calibrated model for the region (Lollato et al., 2020b). We explored the association of 13 environmental descriptors (Supplementary Table 3) and mean environmental yield (e_j_, Equation 2) using regression tree with *rpart* function available in R (Therneau and Atkinson, 1999). We selected hyperparameters by grid search and retained the simplest model that maximized the R^2^ and minimized the root mean squared error.

### Genotypic data

A panel of 234 wheat cultivars (Supplementary Table 2) were genotyped by genotyping-by-sequencing (GBS). The cultivars were obtained directly from private and public breeding programs, USDA-ARS National Plant Germplasm System or USDA Regional Performance Nursery Program. DNA was isolated from leaf tissues collected at three leaf stage using hexadecyltrimethylammonium bromide (CTAB) method (Doyle and Doyle, 1987), and GBS libraries were prepared using *Pst*I and *Msp*I restriction enzymes (Poland et al., 2012). Size selected libraries were sequenced using NextSeq2000 sequencer (Illumina, San Diego) at the USDA Central Small Grain Genotyping Lab, Manhattan, Kansas. Single nucleotide polymorphisms (SNPs) were called using the Chinese Spring (CS) reference genome RefSeq v2.1 (Zhu et al., 2021) developed by International Wheat Genome Sequencing Consortium (IWGSC), which generated 117,391 SNPs. The SNPs were filtered by retaining those with >5% minor allele frequency (MAF), <20% missing data and <5% heterozygous calls. After filtering, 15,224 high-quality SNPs were imputed using K-nearest neighbors imputation algorithm available within TASSEL v5.0 software (Bradbury et al., 2007).

Genome-wide-association study (GWAS) was conducted to explore the associations between 15,224 SNPs and four yield phenotypes (Equations 2, 5) using a multi-locus, fixed and random model circulating probability unification (FarmCPU) available within the GAPIT package in R (Wang and Zhang, 2021). The model was implemented by testing three to 20 principal components (PCs) as covariates to account for population structure. Eigenvectors were obtained from principal component analysis of the genotypic matrix. The most parsimonious model was selected based on visual inspection of quantile-quantile plots between predicted and observed *p*-values. The final model included 10 PCs as covariates plus one SNP in chromosome 2A detected to have strong influence capturing structure of the population analyzed. SNP-trait associations were explored using a threshold of −□log10(P)□=□3.0, where ‘P’ represents the *p*-value of the association of a given SNP with the trait of interest.

We explored the allelic frequency changes with year of cultivar release using a sliding window of 10 years starting in 1989 until 2022. For each SNP, we calculated the effect size of homozygous alleles on the four yield traits using linear regressions. Effect sizes were estimated as the difference between favorable and unfavorable alleles for each SNP × trait combination.

## Results

### Rainfall and photothermal quotient associated with yield

Mean cultivar yield averaged 3.4 Mg ha^-1^ and ranged from 2.5 Mg ha^-1^ to 4.0 Mg ha^-1^ (Figure 2A). Variation in yield between genotypes ranged from 0.4 to 2.5 Mg ha^-1^ in low yielding environments and from 4.0 to 8.4 Mg ha^-1^ in high yielding environments (Figure 2A). Mean environmental yield averaged 3.5 Mg ha^-1^, ranged from 0.7 to 8.8 Mg ha^-1^ (Figure 2B) and correlated with rainfall in both the critical period and grain filling, and with the photothermal quotient in the critical period (Figure 2C, 2D). Regression tree including rainfall and photothermal quotient explained 20% of the variance in environmental mean yield with a root mean squared error of 1.1 Mg ha^-1^ (Figure 2C).

### Genotype-dependent phenotypic plasticity of yield is agronomically adaptive in the Central Plains

Yield plasticity varied 2-fold, from 0.66 in cultivar ‘2163’ to 1.32 in ‘LCS Revere’ (Figure 3A, Supplementary Table 2). Plasticity calculated as slope of reaction norms correlated with plasticity estimated as a ratio of variances (Supplementary Figure 2). Yield plasticity was agronomically adaptive, *i.e.,* phenotypes with higher plasticity had substantially higher yield in high yielding environments, captured in a rate of yield increase of 3.4 Mg ha^-1^ per unit plasticity, and slightly higher yield in low yielding environments, with a rate of 0.3 Mg ha^-1^ per unit plasticity (Figure 3B).

**Figure 3.**
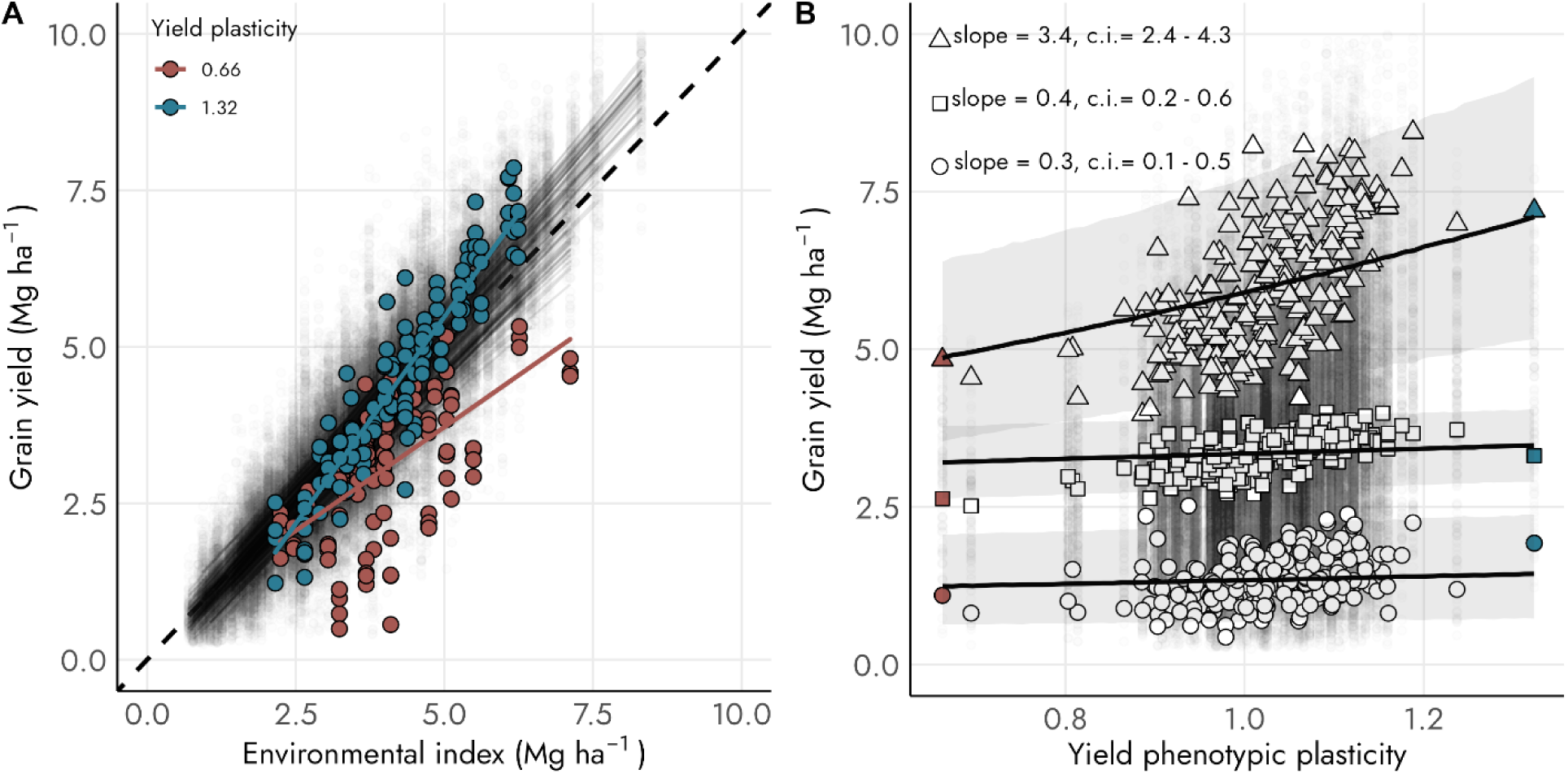
(A) Reaction norms relating genotype-dependent yield and environmental index and (B) association of yield and yield plasticity for 234 cultivars grown in at least 25 environments in Central Great Plains. In (A) color lines and symbols are for cultivars with the highest (blue) and lowest (red) yield plasticity, and black lines are the population as a bundle of reaction norms (*sensu* Van Noordwijk (1989); dashed line is y = x. In (B) triangles are high (95^th^ percentile), squares are average, and circles are low (5^th^ percentile) yielding environments. Model used for quantile regression is described in Equation 5. Environmental index is estimated from the sum of *µ*_0_ and *e_j_* (Equation 2). Yield phenotypic plasticity was estimated as 1+ *β_i_* (Equation 2), *i.e.,* the slope of the reaction norm in panel A.

### Phenotypic plasticity of yield increased and heritability of yield decreased over decades

Yield plasticity increased at 0.5 % year^-1^ (c.i.: 0.4 – 0.7 % year^-1^, Figure 4A). The cultivar with the lowest yield plasticity, ‘2163’, was released in 1989 and its counterpart with highest plasticity, ‘LCS Revere’, in 2018 (Figure 4A). Heritability of yield declined from 0.23 in the decade centered on 1993 to 0.15 in the decade centered on 2017 (Figure 4B).

**Figure 4.**
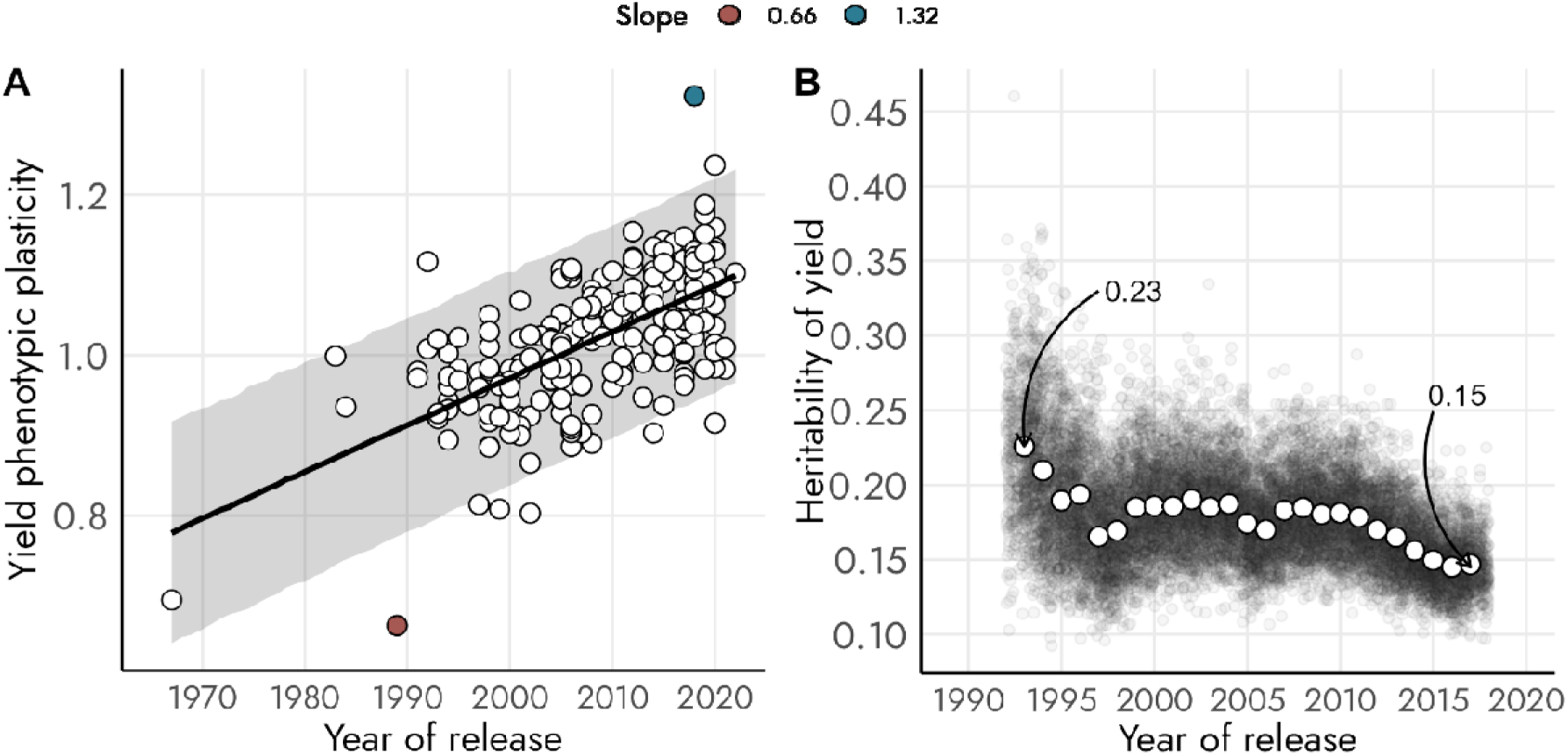
Time trends for (A) phenotypic plasticity of yield, (B) broad sense heritability of yield. In (A), colors show the cultivar with the highest (blue) and lowest (red) yield plasticity. Heritability of yield was estimated using a sliding window of 10 years starting in 1989 until 2022, x-coordinate of open points represents the average heritability for the window sampled. Jitter was added to black points on x, which represent the uncertainty in the estimation of y. In (B), data prior 1989, was excluded as only three cultivars were representative of the entire population in that period.

### Partial independence in the genetic modulation of yield phenotypes

The 2N*^v^*S translocation from *Aegilops ventricosa* Tausch captured population structure in the genotypic data (Figure 5A). The frequency of the 2N*^v^*S translocation in the germplasm pool increased with cultivar year of release, from 6% in 1994 when ‘Jagger’ was released to 68% in 2017 (Figure 5B). Presence of the 2N*^v^*S translocation associated positively with all four yield traits (Figure 5C-F). Presence of the 2N*^v^*S translocation associated with yield more strongly in higher than in lower yielding environments (Figure 5 D vs F, H vs J).

**Figure 5.**
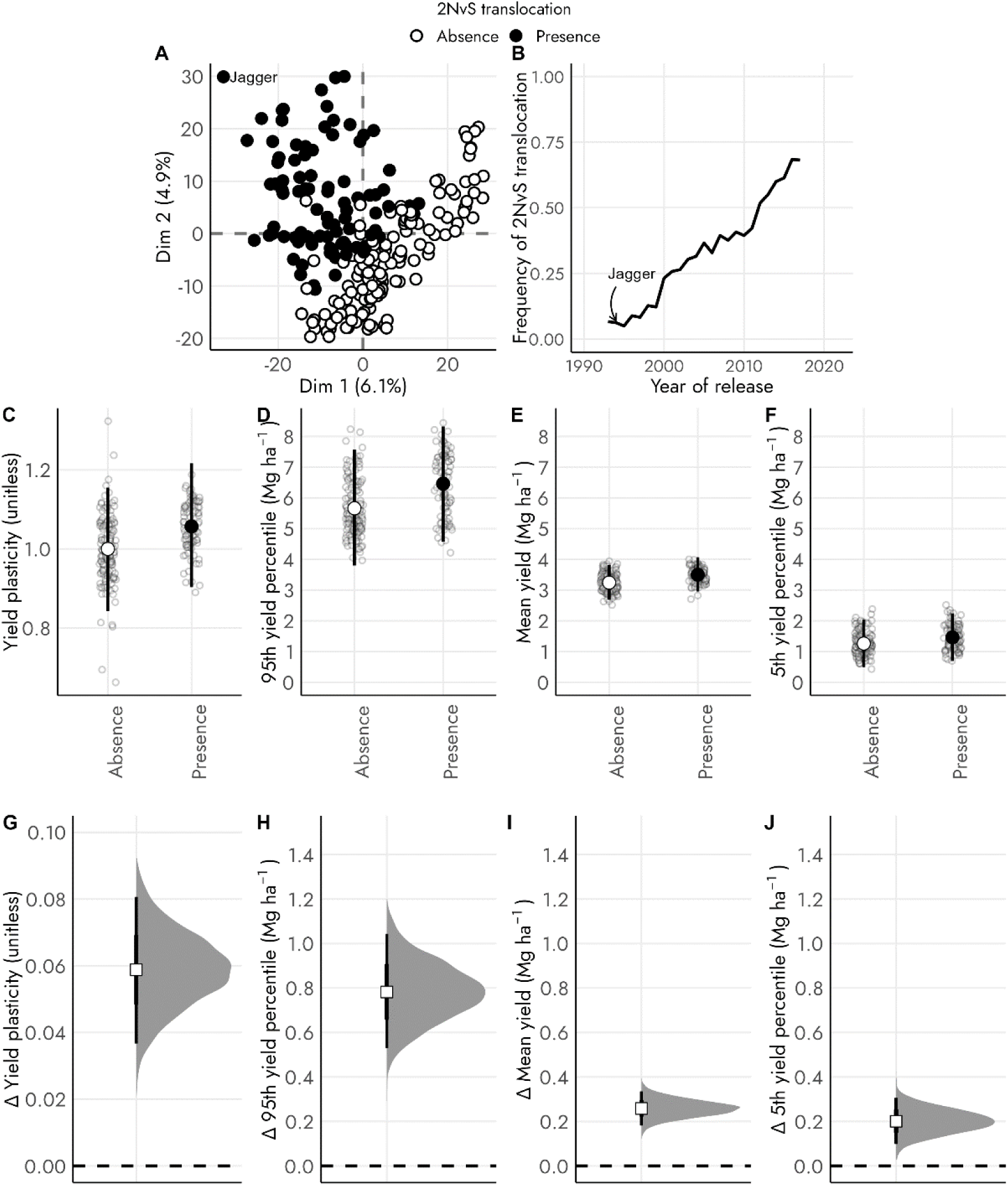
(A) Relationship between first two principal components from PCA conducted on the genotypic matrix. Closed symbols represent presence and open symbols represent absence of 2N*^v^*S translocation on chromosome 2A. Jagger is the canonical for the 2N*^v^*S translocation in North American hard winter wheat. (B) Increasing frequency of the 2N*^v^*S translocation with cultivar year of release. Allelic frequency in the population was calculated using a sliding window of 10 years starting in 1989 until 2022; data prior 1989 were scarce, and therefore were excluded. (C-F) Distribution of yield phenotypes for cultivars with presence and absence of the 2N*^v^*S translocation, circles represent the median and whiskers the 95% credible interval of the posterior predictive distribution of yield phenotypes, while open points represent the observed yield phenotypes. (G-J) Posterior distribution of the difference between presence and absence of the 2N*^v^*S translocation, showing magnitude of effect sizes for each yield phenotype, points represent the mean and whiskers the 95% credible interval. 95% credible interval does not overlap with zero in all four comparisons shown in panels F-I.

We found 46 SNPs above the exploratory threshold of −log_10_(P) = 3.0 to be associated with yield traits including 8 associated with phenotypic plasticity of yield, 12 associated with 95^th^ yield percentile, 12 associated with mean yield, and 14 with 5^th^ yield percentile (Figure 6A-D).

**Figure 6.**
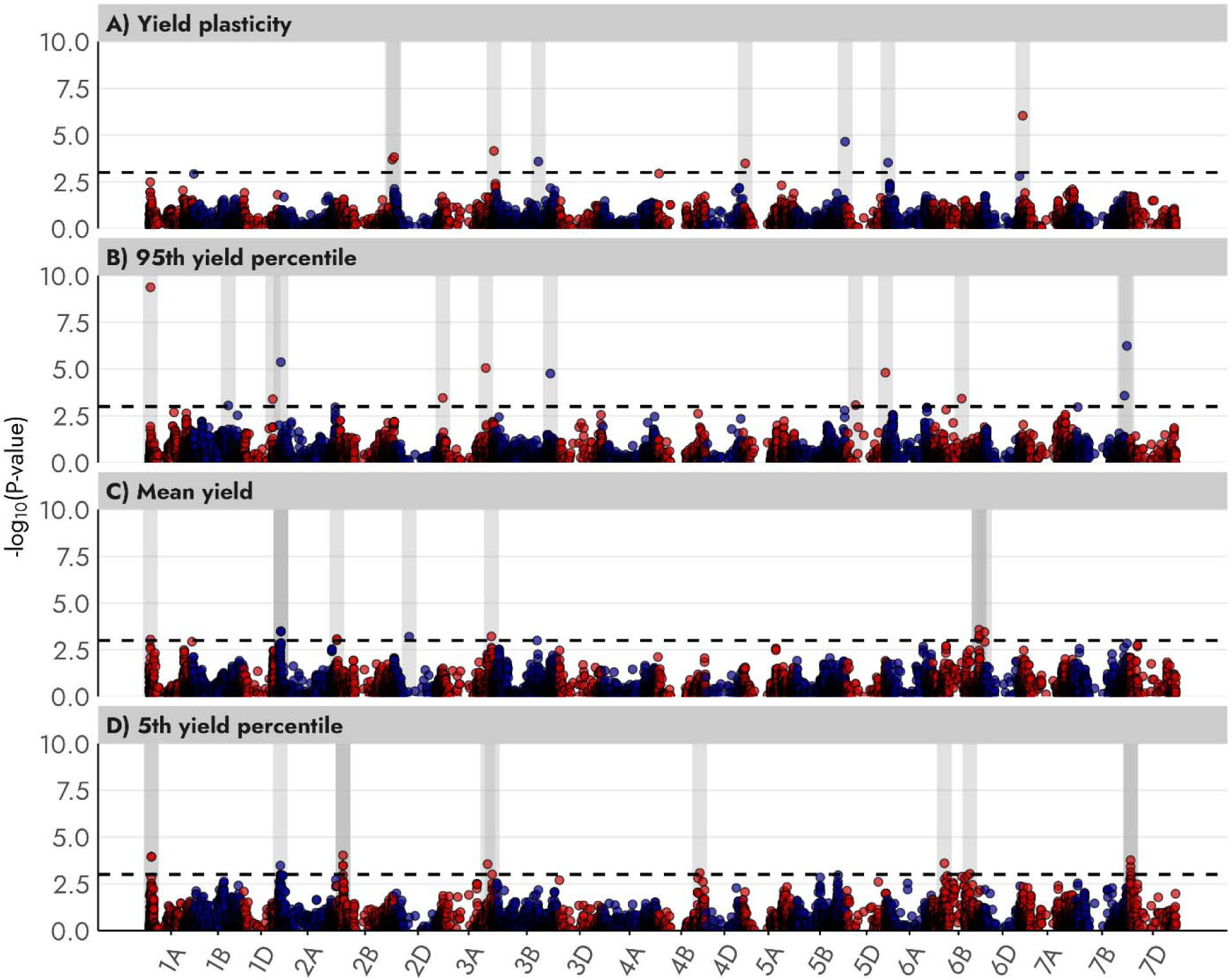
Associations of 15,224 SNPs and four yield phenotypes: (A) phenotypic plasticity of yield, (B) yield in high yielding environments (95^th^ yield percentile), (C) mean yield, and (D) yield in low yielding environments (5^th^ yield percentile). Black dashed line is the exploratory threshold of −log_10_(P) = 3.0, and shaded areas show the position of SNPs above the exploratory threshold.

Frequency of favorable alleles contributing to yield phenotypes changed over time (Figure 7A-D). Increasing yield plasticity with time of cultivar release (Figure 4A) could be partially associated with higher frequency of favorable alleles at SNPs S6A_13708715, S2B_781222236 (Figure 7A). Three SNPs (S6B_412147528, S5D_134802435, S1A_14289872) associated with yield in high yield environments had higher frequency of favorable alleles in newer cultivars (Figure 7B). The frequency of the favorable alleles associated with mean yield, SNPs S2B_13713678, S3A_691148254 and S2D_206645004, increased from older to newer cultivars (Figure 7C). The frequency of favorable alleles associated with yield in low yield environments, SNPs S2A_17606976 and S7D_13070614, increased with time of cultivars release (Figure 7D).

**Figure 7.**
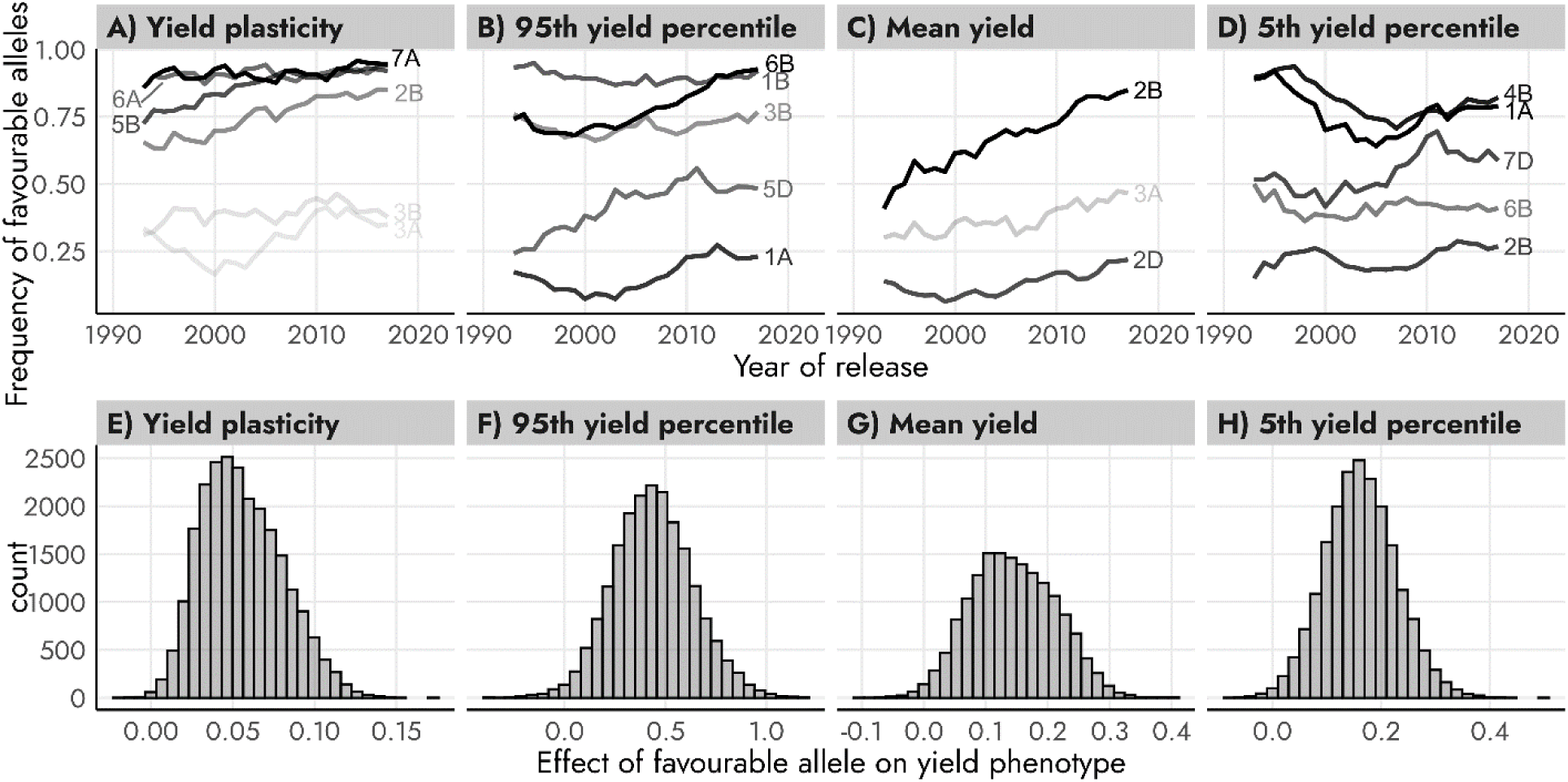
(A-D) Variation in the frequency of the favorable alleles associated with four yield phenotypes with cultivar year of release. Allelic frequency in the population was calculated using a sliding window of 10 years starting in 1989 until 2022; data prior 1989 were scarce, and therefore were excluded. Opacity of the allelic frequency lines is relative effect of the favorable allele in each SNP, being the darker colors the SNPs with major effects on traits. (E-H) Distribution of allelic effects for yield traits. Yield plasticity is unitless in E, and units are Mg ha^-1^ for yield phenotypes in F, G and H. Relative effects represent the effect size of a given SNP relative to the maximum effect size for a given trait.

## Discussion

### Agronomically adaptive changes on wheat yield phenotypes

‘The environment can be source of evolutionary novelties (p.503 Ch. 26, in West-Eberhard, 2003a)’.

West-Eberhard defined evolution as ‘phenotypic change *involving* gene frequency change’ (p.28 Ch. 2, in West-Eberhard, 2003). Following this definition, from a developmental perspective, adaptive evolution occurs when phenotypic variation, triggered by development, is followed by selection over genetically distinct individuals. Our study showed that increasing phenotypic plasticity of yield and changes in favorable allele frequency associated with yield plasticity were a consequence of wheat breeding that drives agronomically adaptive changes. The outcome is that newer cultivars have better adaptation to high rainfall-high photothermal quotient scenarios and cope better with drought stress.

From an agronomic perspective, cultivars with high yield plasticity allow for greater impacts of enhanced agronomy on yield in high yielding environments (Jaenisch et al., 2019; Raj et al., 2023; Giordano et al., 2024). In the current study, we cannot attribute yield gains entirely to higher phenotypic plasticity of recent cultivars. To support the synergy between genetics and agronomy underlying yield gains in farmers’ fields, we introduce three examples of experiments conducted in the region with a sample of cultivars evaluated in this study. Jaenisch et al., (2019) evaluated wheat yield response to a factorial combination of six agronomic practices including nitrogen, sulfur, chloride, foliar fungicides and plant growth regulators and found that management intensification increased environmental mean yield; Raj et al., (2023) reported a divergent bundle of reaction norms of yield with increasing environmental mean yield when the source of variation was management intensity; Giordano et al., (2024) found that cultivars with high yield plasticity outyielded their counterparts with lower plasticity under high yield non-nitrogen limiting environments. Our findings in the current study reinforce associations where, i) enhanced agronomic management improves yield in high yielding environments, ii) a stronger association between yield and yield plasticity in higher yielding environments (Figures 1, 3) and consequently iii) selection for cultivars with higher yield plasticity favors response to enhanced agronomic management.

From a breeding perspective, response to selection (*i.e.,* genetic gain) for a given trait depends on the intensity of selection, the trait heritability, and genetic and phenotypic variance for the trait in the population (Lush, 1937; Cooper and Messina, 2023). A more recent formulation of the genetic gain equation, includes terms of selection accuracy (Cooper and Messina, 2023), which could be the one most impacted when making selections in high yielding environments, where yield differences among cultivars with contrasting plasticity are more evident (Figure 3B). Recent evidence suggested wheat genetic gains are decelerating in the US hard winter wheat region (Maeoka et al., 2020), which might be partially attributed to declining in heritability (Figure 4B), and increasing environmental variability in the context of climate change (Asseng et al., 2011; Tack et al., 2015; Zhao et al., 2017; Zhang et al., 2024).

### Genetic modulation of yield phenotypes

There are bi-directional relationships between the phenotype and both the genome and gene expressivity (West-Eberhard, 2003b; Noble, 2010; Baverstock, 2021; Shapiro, 2022). Here we show that the associations of SNPs with yield varied between favorable and droughted environments, and that plasticity has its own partially independent genetic modulation (Figure 6), aligning with Bradshaw’s (1965) insightful proposition. Empirical evidence from 5,000 *Zea mays* recombinant inbred lines, showed that for 23 agronomically relevant traits, trait means and their plasticities have partially independent genetic modulation and associated candidate genes forming structurally and functionally distinct groups (Kusmec et al., 2017). Reymond et al. (2003) established reaction norms relating maize leaf expansion rate and meristem temperature, air vapor pressure deficit and predawn leaf water potential, showing that the intercept and slope of the reaction norms associated with different QTL. With this approach, they pre-empted QTL x E interaction: instead of asking for QTL associated with leaf expansion rate, and then dealing with the interaction, they asked what QTL associate with the response of the trait to the environment. This approach complements statistical methods to untangle G x E (*e.g.*, Gauch Jr. et al., 2011) but is biologically more insightful. Likewise, we ask: which genomic regions associate with phenotypic plasticity of yield, effectively pre-empting the G x E component of yield variation (Figure 6A).

The 2N*^v^*S translocation from *Aegilops ventricosa* captured most of the population structure on the genome-wide-association analysis (Figure 5). This translocation was first introduced to commercial breeding programs through cultivar ‘Jagger’, released in 1994. The frequency of the 2N*^v^*S in the germplasm pool in this study increased over time (Figure 5B). The 2N*^v^*S translocation has 535 high-confidence genes (Gao et al., 2021) some found to be associated with resistance to biotrophic parasites such as root knot nematodes (*Meloidogyne spp.,* Williamson et al., 2013) and fungal diseases including leaf rust (caused by *Puccinia triticina,* Xue et al., 2018), stripe rust (caused by *P. striiformis f. sp. tritici,* Fang et al., 2011; Mustahsan et al., 2023), stem rust (caused by *P. graminis f. sp. tritici,* Turner et al., 2016) and wheat blast (caused by the *Magnaporthe oryzae triticum,* Cruz et al., 2016; Cruppe et al., 2021). Gao et al., (2021) found that the presence of the 2N*^v^*S translocation was associated positively with yield: i) in 17 out of 19 years of trials in a set of advanced breeding lines selected in the Central Great Plains and ii) in cultivars with CIMMYT germplasm evaluated under irrigation and low disease incidence in Ciudad Obregon (Mexico). Aligning with Gao et al. (2021), our study demonstrated that the 2N*^v^*S translocation was associated with the population structure, suggesting that other genetic variants might be associated with the four yield phenotypes besides the 2N*^v^*S from Jagger, as a major parent used in the regional breeding programs.

#### Box 1

##### Genes associated with yield phenotypes

We tested for the association of SNPs with -□log10(P)□> 3 and annotated genes within the IWGSC CS RefSeq v2.1 (Zhu et al., 2021) to identify putative candidate genes underlying the four yield phenotypes. We set a window size to 20kb centered on each significant SNP.

*TraesCS3A03G1184000* associated with yield plasticity, encodes E3 ubiquitin-protein ligase gene. These proteins are well-known for their role in regulating signaling pathways involving jasmonate, salicylic acid, ethylene, gibberellic acid, and auxin in response to various stresses (Vierstra, 2012; Shabek and Zheng, 2014; Kelley, 2018). The specific functions of E3 ubiquitin ligase proteins in response to environmental biotic and abiotic stresses have been characterized in wheat, soybeans and rice (Su et al., 2024). In wheat, the E3 ubiquitin gene *CMPG1–V* overexpression was associated with increased resistance to powdery mildew during both the seedling and adult stages (Zhu et al., 2021). E3 ubiquitin proteins from wheat may either positively or negatively regulate response to salt and drought stress (Zhang et al., 2017; Wu et al., 2020). *TraesCS2B03G1463700* encodes a Protein S-acyltransferase (PAT) 11 gene.

*Arabidopsis* mutants lacking *PAT14* gene had premature leaf senescence (Zhao et al., 2016). *TraesCS6A03G0054100* associated with yield plasticity and *TraesCS3B03G1223300* associated with yield in high yielding environments (Box 1 Table). These two genes were situated near F-box proteins. This protein family, was identified in rice (Kuroda et al., 2002; Hua et al., 2011), maize (Jia et al., 2013), soybeans (Jia et al., 2017) and *Arabidopsis* (Hua et al., 2011). A major study on F-box genes in wheat identified 409 genes spread across all 21 chromosomes (Li et al., 2020), and were expressed during the formation of spikes, grains, and flag leaves. F-box genes associated with resistance to leaf rust, drought, heat and cold stress (Li et al., 2020). The F-box protein gene, *TaCFBD*, cloned from wheat, was overexpressed when wheat seedlings were exposed to a temperature of 4°C and overexpressed during the early stages of inflorescence development, suggesting a potential connection to the formation of floral organs (Hong et al., 2012).

Three characterized genes associated with yield in high yielding environments (Box 1 Table). *TraesCS5D03G1145000,* encodes a Phospholipase D (PDL), involved in tolerance against freezing, dehydration, heat, and salt stress in wheat, *Arabidopsis*, maize, sorghum and potato (Wang et al., 2021; Wei et al., 2022). *TraesCS7B03G1226200* encodes a GDSL lipase/esterase protein (Glycine – Aspartic acid – Serine – Leucine). GDSL lipase/esterase (GELP) were identified in *Arabidopsis*, maize, soybean and rice (Shen et al., 2022) and are involved in seed germination, coleoptile elongation, lateral root formation, overall root development, plant height, and development of stomata and reproductive structures (Ding et al., 2019). 509 GELP genes were identified in wheat, *TaGELP073*, has a role in the development of anthers and pollen (Yang et al., 2022).

Five characterized candidate genes associated with mean yield (Box 1 Table). *TraesCS2D03G0491900* encodes a helicase-conserved C-terminal domain protein. DNA helicases are involved in genetic information processing, including replication, transcription, translation, repair, and recombination (Umate et al., 2010). A genome-wide analysis of an RNA helicase family in wheat showed that certain genes within this family were upregulated in transgenic *Arabidopsis* plants (with ectopic expression of specific wheat helicase genes) under drought, salt, and cold stress conditions (Ru et al., 2021). *TraesCS2D03G0492100* encodes MAIN-LIKE 1 protein. The MAINTENANCE OF MERISTEMS (MAIN) genes play a crucial role in maintaining genome stability, developmental processes, and silencing transposable elements (Nicolau et al., 2020). In *Arabidopsis,* mutations in the MAIN gene result in premature differentiation and cell death of stem cells, shorter roots, irregularly shaped leaves, reduced fertility, and partial fasciation of stems (Wenig et al., 2013). *TraesCS6B03G1046500* encodes subtilisin-like serine protease (SBT), a subgroup of the serine protease family. In wheat, 255 putative subtilisin-like serine protease (SBT) genes were identified (Zhao et al., 2023) and expression levels of SBT genes varied with fungal and bacterial pathogens, as well as under conditions of drought, heat, and cold, expressed during seedling emergence, tillering, and grain filling period (Zhao et al., 2023). In cotton (*Gossypium hirsutum*), it is hypothesized that the SBT gene *GhSBT27A* enhances drought tolerance by altering the structure of chloroplasts (Dai et al., 2023). *TraesCS6B03G1290400* inhibits pectin methylesterase, its expression in wheat fertile anthers, suggests a potential role in pollen development and fertility (Wang et al., 2023).

Yield in low yielding environments was primarily a response to drought stress, and associated with six characterized genes (Box 1 Table). *TraesCS7D03G0060000*, encodes late embryogenesis abundant (LEA) proteins, its expression in wheat roots, stems, leaves, and seeds varied with environment and associated with drought stress and low-temperature stress (Zan et al., 2020). *TraesCS1A03G0106400* encodes ankyrin repeat (ANK) proteins, which modulate abscisic acid signaling pathways and are associated with tiller number (Dong et al., 2023) and resistance to various fungal diseases, including leaf rust (Kolodziej et al., 2021), stripe rust (Wang et al., 2020), and powdery mildew (Hu et al., 2022). *TraesCS7D03G0060600* encodes a putative β-1,3 glucan synthase (callose synthase 10) gene. Callose plays a crucial role in the transmission of information between cells and regulates growth and development at various stages (Li et al., 2023). Both biotic and abiotic stresses can stimulate the synthesis of callose in plants (Li et al., 2023). Bacterial infections can trigger callose synthesis at the location of the attack, resulting in thicker cell walls forming a protective physical barrier that aids the plant in defending itself against additional bacterial intrusion (Li et al., 2023).

**Box 1 Table.** Genes associated with yield phenotypes in a 20kb window centered on each SNP above the −log10(P) = 3.0 threshold. Genes were described using the Chinese Spring (CS) reference genome RefSeq v2.1 (Zhu et al., 2021). Abbreviations: chromosome (Chr.), base pairs (bp).

| Trait | Chr. | Physical location (bp) | Gene Name | Description |
| --- | --- | --- | --- | --- |
| Yield plasticity | 2B | 781222236 | *TraesCS2B03G1463700* | Protein S-acyltransferase 11 |
|  | 3A | 726890110 | *TraesCS3A03G1184000* | E3 ubiquitin-protein ligase RGLG3 |
|  | 3B | 587824478 | *TraesCS3B03G0914100* | Uncharacterized protein |
|  | 3B | 587824478 | *TraesCS3B03G0914200* | Basic-leucine zipper domain; |
|  | 5B | 704099016 | *TraesCS5B03G1336800* | Uncharacterized protein |
|  | 6A | 13708715 | *TraesCS6A03G0054100* | F-box domain |
| 95th yield percentile | 1B | 493165543 | *TraesCS1B03G0773600* | Uncharacterized protein |
|  | 3B | 754137029 | *TraesCS3B03G1223300* | F-box domain |
|  | 5D | 134802435 | *TraesCS5D03G0270200* | Uncharacterized protein |
|  | 5D | 547030857 | *TraesCS5D03G1145000* | Phospholipase D delta |
|  | 5D | 547030857 | *TraesCS5D03G1145100* | Uncharacterized protein |
|  | 7B | 724961593 | *TraesCS7B03G1226200* | GDSL lipase/esterase |
| Mean yield | 2D | 206645004 | *TraesCS2D03G0491900* | Helicase conserved C-terminal domain |
|  | 2D | 206645004 | *TraesCS2D03G0492100* | Protein MAIN-LIKE 1 |
|  | 6B | 649190074 | *TraesCS6B03G1038400* | Serine-threonine/tyrosine-protein kinase, catalytic domain RIPK |
|  | 6B | 651828250 | TraesCS6B03G1046400 | Uncharacterized protein |
|  | 6B | 651828250 | TraesCS6B03G1046500 | Subtilisin-like protease SBT1.4 |
|  | 6B | 726368246 | TraesCS6B03G1290400 | Pectinesterase inhibitor domain |
|  | 6B | 726368246 | TraesCS6B03G1290500 | Uncharacterized protein |
| 5th yield percentile | 1A | 27594591 | TraesCS1A03G0106400 | Ankyrin repeats |
|  | 1A | 27594593 | TraesCS1A03G0106400 | Ankyrin repeats |
|  | 3A | 701725274 | TraesCS3A03G1092300 | Uncharacterized protein |
|  | 7D | 13070614 | TraesCS7D03G0060000 | Late embryogenesis abundant protein, LEA-25/LEA-D113 |
|  | 7D | 13070644 | TraesCS7D03G0060000 | Late embryogenesis abundant protein, LEA-25/LEA-D113 |
|  | 7D | 13415400 | TraesCS7D03G0060600 | 1,3-beta-glucan synthase subunit FKS1, domain-1/Callose synthase 10 |

## Conclusion

In the context of large investments in genome-wide-association studies of quantitative traits including yield and its components (Zhang et al., 2010; Quraishi et al., 2017; Yang et al., 2021), here we show the mismatch between genomic regions associated with mean yield, the most common target of GWAS, yield in high and low yielding environments, and between yield and phenotypic plasticity of yield. Our findings have implications for research, breeding and agronomy. A focus on phenotypic plasticity could be useful to pre-empt GxE. First, to exploit the genetics of yield plasticity, we need to determine whether yield plasticity is agronomically adaptive for the population of genotypes and environments evaluated. Further exploration of data from regional testing networks in major wheat growing regions could be used to answer this context-specific question. Second, selection for yield plasticity may improve selection accuracy and intensity, arising as a potential tool to overcome lower genetic gains partially associated with declining heritability of yield. Third, selection on high yielding environments allows to better detect differences in cultivar-specific phenotypic plasticity. However, maintaining a necessary level of testing in lower yielding environments will capture genomic regions associated with stress adaptation. Fourth, breeders can select semi-independently for yield potential, drought stress tolerance and yield plasticity. Finally, a dual focus on phenotypes with high yield plasticity and enhanced management could further capture breeding and agronomy synergies.

## Acknowledgements

Mention of trade names or commercial products in this publication is solely for the purpose of providing specific information and does not imply recommendation or endorsement by the U.S. Department of Agriculture. USDA is an equal opportunity provider and employer. The authors acknowledge support received from Edward Souza and members of the Alberta Regional Variety Advisory Committee and Manitoba Crop Variety Evaluation Team in helping to collect data used in this manuscript. We also acknowledge support received from Skyla Jost on laboratory procedures for DNA data collection and support from Eric Fabrizius who gave us seeds from much of the germplasms used in this study. This is research contribution no. 25-081-J from the Kansas Agricultural Experiment Station.

## Author contributions

NG, VOS, MJG, RL : conceptualization; NG, VOS, MJG, RL: method; NG, GC: formal analysis; NG: writing - original draft; NG: visualization; NG, AKF, GB, RL: funding acquisition; NG, VOS, MJG, RL, GC, GB, AKF, SH, GZ, JB: writing - review & editing; PA, AB, AOS, SJD, JL, SH, GZ: investigation, data curation; GB,AB, PSA: investigation, resources; JB: resources.

## Conflict of interest

The authors declare no conflicts of interest.

## Funding

This work was partially funded by Kansas Wheat Alliance.

## Data availability

Restrictions apply to the availability of data shown in Figure 1, which were used under license for this study. Data from Figure 1 can be requested directly from the institutions who shared the data. All other data used in this study are available upon request to the authors. Scripts can be found in https://github.com/giordanon/Yield-Plasticity-GWAS.

## Abbreviations

c.i.: Credible interval
G × E: genotype × environment
CIMMYT: Centro Internacional de Mejoramiento de Maiz y Trigo
SNPs: single nucleotide polymorphisms
GWAS: genome-wide-association study

## Notes

### Competing Interest Statement

The authors have declared no competing interest.

## References

Alvarez Prado, S., V.O. Sadras, and L. Borrás. 2014. Independent genetic control of maize (Zea mays L.) kernel weight determination and its phenotypic plasticity. Journal of Experimental Botany 65(15): 4479–4487. doi: 10.1093/jxb/eru215.

Amzallag, G.N. 2000. Connectance in *Sorghum* development: beyond the genotype–phenotype duality. Biosystems 56(1): 1–11. doi: 10.1016/S0303-2647(00)00068-X.

Asseng, S., I. Foster, and N.C. Turner. 2011. The impact of temperature variability on wheat yields. Global Change Biology 17(2): 997–1012. doi: 10.1111/j.1365-2486.2010.02262.x.

Auld, J.R., A.A. Agrawal, and R.A. Relyea. 2009. Re-evaluating the costs and limits of adaptive phenotypic plasticity. Proceedings of the Royal Society B: Biological Sciences 277(1681): 503–511. doi: 10.1098/rspb.2009.1355.

Battenfield, S.D., A.R. Klatt, and W.R. Raun. 2013. Genetic Yield Potential Improvement of Semidwarf Winter Wheat in the Great Plains. Crop Science 53(3): 946–955. doi: 10.2135/cropsci2012.03.0158.

Baverstock, K. 2021. The gene: An appraisal. Progress in Biophysics and Molecular Biology 164: 46–62. doi: 10.1016/j.pbiomolbio.2021.04.005.

Berzonsky, W.A., and H.N. Lafever. 1993. Progress in Ohio Soft Red Winter Wheat Breeding: Grain Yield and Agronomic Traits of Cultivars Released from 1871 to 1987. Crop Science 33(6): cropsci1993.0011183X003300060050x. doi: 10.2135/cropsci1993.0011183X003300060050x.

Bradbury, P.J., Z. Zhang, D.E. Kroon, T.M. Casstevens, Y. Ramdoss, et al. 2007. TASSEL: software for association mapping of complex traits in diverse samples. Bioinformatics 23(19): 2633–2635. doi: 10.1093/bioinformatics/btm308.

Bradshaw, A.D. 1965. Evolutionary Significance of Phenotypic Plasticity in Plants. Advances in Genetics. Elsevier. p. 115–155

Brisson, N., P. Gate, D. Gouache, G. Charmet, F.-X. Oury, et al. 2010. Why are wheat yields stagnating in Europe? A comprehensive data analysis for France. Field Crops Research 119(1): 201–212. doi: 10.1016/j.fcr.2010.07.012.

Cooper, M., and C.D. Messina. 2023. Breeding crops for drought-affected environments and improved climate resilience. The Plant Cell 35(1): 162–186. doi: 10.1093/plcell/koac321.

Cossani, C.M., and V.O. Sadras. 2021. Nitrogen and water supply modulate the effect of elevated temperature on wheat yield. European Journal of Agronomy 124: 126227. doi: 10.1016/j.eja.2020.126227.

Cox, T.S., J.P. Shroyer, L. Ben-Hui, R.G. Sears, and T.J. Martin. 1988. Genetic Improvement in Agronomic Traits of Hard Red Winter Wheat Cultivars 1919 to 1987. Crop Science 28(5): cropsci1988.0011183X002800050006x. doi: 10.2135/cropsci1988.0011183X002800050006x.

Cruppe, G., P. Silva, C. Lemes da Silva, G. Peterson, K.F. Pedley, et al. 2021. Genome-wide association reveals limited benefits of pyramiding the 1B and 1D loci with the 2NvS translocation for wheat blast control. Crop Science 61(2): 1089–1103. doi: 10.1002/csc2.20397.

Cruz, C. d., G. l. Peterson, W. w. Bockus, P. Kankanala, J. Dubcovsky, et al. 2016. The 2NS Translocation from Aegilops ventricosa Confers Resistance to the Triticum Pathotype of Magnaporthe oryzae. Crop Science 56(3): 990–1000. doi: 10.2135/cropsci2015.07.0410.

Dai, M., N. Zhou, Y. Zhang, Y. Zhang, K. Ni, et al. 2023. Genome-wide analysis of the SBT gene family involved in drought tolerance in cotton. Front. Plant Sci. 13: 1097732. doi: 10.3389/fpls.2022.1097732.

Dewitt, T., and S. Scheiner. 2004.Phenotypic plasticity: Functional and conceptual approaches.

DeWitt, T.J., A. Sih, and D.S. Wilson. 1998. Costs and limits of phenotypic plasticity. Trends in Ecology & Evolution 13(2): 77–81. doi: 10.1016/S0169-5347(97)01274-3.

Ding, L.-N., M. Li, W.-J. Wang, J. Cao, Z. Wang, et al. 2019. Advances in plant GDSL lipases: from sequences to functional mechanisms. Acta Physiol Plant 41(9): 151. doi: 10.1007/s11738-019-2944-4.

Dingemanse, N.J., A.J.N. Kazem, D. Réale, and J. Wright. 2010. Behavioural reaction norms: animal personality meets individual plasticity. Trends in Ecology & Evolution 25(2): 81–89. doi: 10.1016/j.tree.2009.07.013.

Diouf, I., L. Derivot, S. Koussevitzky, Y. Carretero, F. Bitton, et al. 2020. Genetic basis of phenotypic plasticity and genotype × environment interactions in a multi-parental tomato population. Journal of Experimental Botany 71(18): 5365–5376. doi: 10.1093/jxb/eraa265.

Dong, C., L. Zhang, Q. Zhang, Y. Yang, D. Li, et al. 2023. Tiller Number1 encodes an ankyrin repeat protein that controls tillering in bread wheat. Nat Commun 14(1): 836. doi: 10.1038/s41467-023-36271-z.

Donmez, E., R.G. Sears, J.P. Shroyer, and G.M. Paulsen. 2001. Genetic Gain in Yield Attributes of Winter Wheat in the Great Plains. Crop Science 41(5): 1412–1419. doi: 10.2135/cropsci2001.4151412x.

Donovan, L.A., H. Maherali, C.M. Caruso, H. Huber, and H. de Kroon. 2011. The evolution of the worldwide leaf economics spectrum. Trends in Ecology & Evolution 26(2): 88–95. doi: 10.1016/j.tree.2010.11.011.

Doyle, J.J., and J.L. Doyle, editors. 1987. A rapid DNA isolation procedure for small quantities of fresh leaf tissue. PHYTOCHEMICAL BULLETIN.

Dupont, F.M., W.J. Hurkman, W.H. Vensel, C. Tanaka, K.M. Kothari, et al. 2006. Protein accumulation and composition in wheat grains: Effects of mineral nutrients and high temperature. European Journal of Agronomy 25(2): 96–107. doi: 10.1016/j.eja.2006.04.003.

Fang, T., K.G. Campbell, Z. Liu, X. Chen, A. Wan, et al. 2011. Stripe Rust Resistance in the Wheat Cultivar Jagger is Due to Yr17 and a Novel Resistance Gene. Crop Science 51(6): 2455– 2465. doi: 10.2135/cropsci2011.03.0161.

de Felipe, M., and S. Alvarez Prado. 2021. Has yield plasticity already been exploited by soybean breeding programmes in Argentina? Journal of Experimental Botany 72(20): 7264– 7273. doi: 10.1093/jxb/erab347.

Fischer, R.A. 1985. Number of kernels in wheat crops and the influence of solar radiation and temperature. J. Agric. Sci. 105(2): 447–461. doi: 10.1017/S0021859600056495.

Fischer, T., D. Byerlee, and G. Edmeades. 2014. Crop yields and global food security: will yield increase continue to feed the world? ACIAR, Canberra.

Fischer, R.A., and G.J. Rebetzke. 2018. Indirect selection for potential yield in early-generation, spaced plantings of wheat and other small-grain cereals: a review. Crop Pasture Sci. 69(5): 439. doi: 10.1071/CP17409.

Fufa, H., P.S. Baenziger, B.S. Beecher, R.A. Graybosch, K.M. Eskridge, et al. 2005. Genetic improvement trends in agronomic performances and end-use quality characteristics among hard red winter wheat cultivars in Nebraska. Euphytica 144(1): 187–198. doi: 10.1007/s10681-005-5811-x.

Gao, L., D.-H. Koo, P. Juliana, T. Rife, D. Singh, et al. 2021. The Aegilops ventricosa 2NvS segment in bread wheat: cytology, genomics and breeding. Theor Appl Genet 134(2): 529–542. doi: 10.1007/s00122-020-03712-y.

Gauch Jr., H.G., P.C. Rodrigues, J.D. Munkvold, E.L. Heffner, and M. Sorrells. 2011. Two New Strategies for Detecting and Understanding QTL × Environment Interactions. Crop Science 51(1): 96–113. doi: 10.2135/cropsci2010.04.0206.

Giordano, N., V.O. Sadras, A.A. Correndo, and R.P. Lollato. 2024. Cultivar-specific phenotypic plasticity of yield and grain protein concentration in response to nitrogen in winter wheat. Field Crops Research 306: 109202. doi: 10.1016/j.fcr.2023.109202.

Green, A.J., G. Berger, C.A. Griffey, R. Pitman, W. Thomason, et al. 2012. Genetic Yield Improvement in Soft Red Winter Wheat in the Eastern United States from 1919 to 2009. Crop Science 52(5): 2097–2108. doi: 10.2135/cropsci2012.01.0026.

Grogan, S.M., J. Anderson, P. Stephen Baenziger, K. Frels, M.J. Guttieri, et al. 2016. Phenotypic plasticity of winter wheat heading date and grain yield across the US great plains. Crop Science 56(5): 2223–2236. doi: 10.2135/cropsci2015.06.0357.

Hay, R.K.M. 1995. Harvest index: a review of its use in plant breeding and crop physiology. Annals of Applied Biology 126(1): 197–216. doi: 10.1111/j.1744-7348.1995.tb05015.x.

Hong, M.J., D.Y. Kim, S.Y. Kang, D.S. Kim, J.B. Kim, et al. 2012. Wheat F-box protein recruits proteins and regulates their abundance during wheat spike development. Mol Biol Rep 39(10): 9681–9696. doi: 10.1007/s11033-012-1833-3.

Hu, P., Y. Ren, J. Xu, Q. Wei, P. Song, et al. 2022. Identification of ankyrin-transmembrane-type subfamily genes in Triticeae species reveals TaANKTM2A-5 regulates powdery mildew resistance in wheat. Front. Plant Sci. 13: 943217. doi: 10.3389/fpls.2022.943217.

Hua, Z., C. Zou, S.-H. Shiu, and R.D. Vierstra. 2011. Phylogenetic Comparison of F-Box (FBX) Gene Superfamily within the Plant Kingdom Reveals Divergent Evolutionary Histories Indicative of Genomic Drift. PLOS ONE 6(1): e16219. doi: 10.1371/journal.pone.0016219.

Hufkens, K., D. Basler, T. Milliman, E.K. Melaas, and A.D. Richardson. 2018. An integrated phenology modelling framework in r. Methods in Ecology and Evolution 9(5): 1276–1285. doi: 10.1111/2041-210X.12970.

Jaenisch, B.R., A. de Oliveira Silva, E. DeWolf, D.A. Ruiz-Diaz, and R.P. Lollato. 2019. Plant Population and Fungicide Economically Reduced Winter Wheat Yield Gap in Kansas. Agronomy Journal 111(2): 650–665. doi: 10.2134/agronj2018.03.0223.

Jia, F., B. Wu, H. Li, J. Huang, and C. Zheng. 2013. Genome-wide identification and characterisation of F-box family in maize. Mol Genet Genomics 288(11): 559–577. doi: 10.1007/s00438-013-0769-1.

Jia, Q., Z.-X. Xiao, F.-L. Wong, S. Sun, K.-J. Liang, et al. 2017. Genome-Wide Analyses of the Soybean F-Box Gene Family in Response to Salt Stress. Int J Mol Sci 18(4): 818. doi: 10.3390/ijms18040818.

Kelley, D.R. 2018. E3 Ubiquitin Ligases: Key Regulators of Hormone Signaling in Plants *. Molecular & Cellular Proteomics 17(6): 1047–1054. doi: 10.1074/mcp.MR117.000476.

Khalil, I.H., B.F. Carver, E.G. Krenzer, C.T. MacKown, and G.W. Horn. 2002. Genetic Trends in Winter Wheat Yield and Test Weight under Dual-Purpose and Grain-Only Management Systems. Crop Science 42(3): 710–715. doi: 10.2135/cropsci2002.7100.

Kolodziej, M.C., J. Singla, J. Sánchez-Martín, H. Zbinden, H. Šimková, et al. 2021. A membrane-bound ankyrin repeat protein confers race-specific leaf rust disease resistance in wheat. Nat Commun 12(1): 956. doi: 10.1038/s41467-020-20777-x.

Kuroda, H., N. Takahashi, H. Shimada, M. Seki, K. Shinozaki, et al. 2002. Classification and Expression Analysis of Arabidopsis F-Box-Containing Protein Genes. Plant and Cell Physiology 43(10): 1073–1085. doi: 10.1093/pcp/pcf151.

Kusmec, A., N. de Leon, and P.S. Schnable. 2018. Harnessing Phenotypic Plasticity to Improve Maize Yields. Frontiers in Plant Science 9. https://www.frontiersin.org/articles/10.3389/fpls.2018.01377 (accessed 31 March 2023).

Kusmec, A., S. Srinivasan, D. Nettleton, and P.S. Schnable. 2017. Distinct genetic architectures for phenotype means and plasticities in Zea mays. Nature Plants 3(9): 715–723. doi: 10.1038/s41477-017-0007-7.

Lawrence, M., W. Huber, H. Pagès, P. Aboyoun, M. Carlson, et al. 2013. Software for Computing and Annotating Genomic Ranges. PLOS Computational Biology 9(8): e1003118. doi: 10.1371/journal.pcbi.1003118.

Li, N., Z. Lin, P. Yu, Y. Zeng, S. Du, et al. 2023. The multifarious role of callose and callose synthase in plant development and environment interactions. Front. Plant Sci. 14: 1183402. doi: 10.3389/fpls.2023.1183402.

Li, H., C. Wei, Y. Meng, R. Fan, W. Zhao, et al. 2020. Identification and expression analysis of some wheat F-box subfamilies during plant development and infection by *Puccinia triticina*. Plant Physiology and Biochemistry 155: 535–548. doi: 10.1016/j.plaphy.2020.06.040.

Lian, L., and G. Campos. 2015. FW: An R package for Finlay-Wilkinson Regression That Incorporates Genomic/Pedigree Information and Covariance Structures Between Environments. G3 (Bethesda, Md.) 6. doi: 10.1534/g3.115.026328.

Lo Valvo, P.J., D.J. Miralles, and R.A. Serrago. 2018. Genetic progress in Argentine bread wheat varieties released between 1918 and 2011: Changes in physiological and numerical yield components. Field Crops Research 221: 314–321. doi: 10.1016/j.fcr.2017.08.014.

Lollato, R.P., G.P. Bavia, V. Perin, M. Knapp, E.A. Santos, et al. 2020a. Climate-risk assessment for winter wheat using long-term weather data. Agronomy Journal 112(3): 2132– 2151. doi: 10.1002/agj2.20168.

Lollato, R.P., K. Roozeboom, J.F. Lingenfelser, C.L. da Silva, and G. Sassenrath. 2020b. Soft winter wheat outyields hard winter wheat in a subhumid environment: Weather drivers, yield plasticity, and rates of yield gain. Crop Science 60(3): 1617–1633. doi: 10.1002/csc2.20139.

Lush, J.L. 1937. Animal Breeding Plans. Iowa State College Press, Ames, Iowa.

Maeoka, R.E., V.O. Sadras, I.A. Ciampitti, D.R. Diaz, A.K. Fritz, et al. 2020. Changes in the Phenotype of Winter Wheat Varieties Released Between 1920 and 2016 in Response to In-Furrow Fertilizer: Biomass Allocation, Yield, and Grain Protein Concentration. Frontiers in Plant Science 10. https://www.frontiersin.org/articles/10.3389/fpls.2019.01786 (accessed 24 April 2023).

Munaro, L.B., T.J. Hefley, E. DeWolf, S. Haley, A.K. Fritz, et al. 2020. Exploring long-term variety performance trials to improve environment-specific genotype × management recommendations: A case-study for winter wheat. Field Crops Research 255: 107848. doi: 10.1016/j.fcr.2020.107848.

Mustahsan, W., M.J. Guttieri, R.L. Bowden, K. Garland-Campbell, K. Jordan, et al. 2023. Mapping the quantitative field resistance to stripe rust in a hard winter wheat population “Overley” × “Overland.” Crop Science 63(4): 2050–2066. doi: 10.1002/csc2.20977.

Nicolau, M., P. N, D. J, J.-A. Y, F. S, et al. 2020. The plant mobile domain proteins MAIN and MAIL1 interact with the phosphatase PP7L to regulate gene expression and silence transposable elements in Arabidopsis thaliana. PLoS genetics. doi: 10.1371/journal.pgen.1008324.

Noble, D. 2010. Differential and integral views of genetics in computational systems biology. Interface Focus 1(1): 7–15. doi: 10.1098/rsfs.2010.0444.

van Noordwijk, A.J. 1989. Reaction Norms in Genetical Ecology. BioScience 39(7): 453–458. doi: 10.2307/1311137.

Poland, J., J. Endelman, J. Dawson, J. Rutkoski, S. Wu, et al. 2012. Genomic Selection in Wheat Breeding using Genotyping-by-Sequencing. The Plant Genome 5(3). doi: 10.3835/plantgenome2012.06.0006.

Quraishi, U.M., C. Pont, Q. Ain, R. Flores, L. Burlot, et al. 2017. Combined Genomic and Genetic Data Integration of Major Agronomical Traits in Bread Wheat (Triticum aestivum L.). Front. Plant Sci. 8. doi: 10.3389/fpls.2017.01843.

R Core Team. 2021. A language and environment for statistical computing. https://www.R-project.org/.

Raj, A.S., K. Siliveru, R. McLean, P.V.V. Prasad, and R.P. Lollato. 2023. Intensive management simultaneously reduces yield gaps and improves milling and baking properties of bread wheat. Crop Science 63(2): 936–955. doi: 10.1002/csc2.20906.

Reymond, M., B. Muller, A. Leonardi, A. Charcosset, and F. Tardieu. 2003. Combining Quantitative Trait Loci Analysis and an Ecophysiological Model to Analyze the Genetic Variability of the Responses of Maize Leaf Growth to Temperature and Water Deficit. Plant Physiology 131(2): 664–675. doi: 10.1104/pp.013839.

Ru, J.-N., Z.-H. Hou, L. Zheng, Q. Zhao, F.-Z. Wang, et al. 2021. Genome-Wide Analysis of DEAD-box RNA Helicase Family in Wheat (Triticum aestivum) and Functional Identification of TaDEAD-box57 in Abiotic Stress Responses. Front. Plant Sci. 12: 797276. doi: 10.3389/fpls.2021.797276.

Ruiz, M.B., K.E. D’Andrea, and M.E. Otegui. 2019. Phenotypic plasticity of maize grain yield and related secondary traits: Differences between inbreds and hybrids in response to contrasting water and nitrogen regimes. Field Crops Research 239: 19–29. doi: 10.1016/j.fcr.2019.04.004.

Sadras, V.O. 2024. Bread and hummus: trait connectance and correlation pleiades in grain crops. Journal of Experimental Botany: erae374. doi: 10.1093/jxb/erae374.

Sadras, V.O., and C. Lawson. 2011. Genetic gain in yield and associated changes in phenotype, trait plasticity and competitive ability of South Australian wheat varieties released between 1958 and 2007. Crop Pasture Sci. 62(7): 533. doi: 10.1071/CP11060.

Sadras, V.O., and R.A. Richards. 2014. Improvement of crop yield in dry environments: benchmarks, levels of organisation and the role of nitrogen. Journal of Experimental Botany 65(8): 1981–1995. doi: 10.1093/jxb/eru061.

Sadras, V.O., and G.A. Slafer. 2012. Environmental modulation of yield components in cereals: Heritabilities reveal a hierarchy of phenotypic plasticities. Field Crops Research 127: 215–224. doi: 10.1016/j.fcr.2011.11.014.

Scheiner, S.M. 1993. Genetics and Evolution of Phenotypic Plasticity. Annual Review of Ecology, Evolution, and Systematics 24(Volume 24, 1993): 35–68. doi: 10.1146/annurev.es.24.110193.000343.

Shabek, N., and N. Zheng. 2014. Plant ubiquitin ligases as signaling hubs. Nat Struct Mol Biol 21(4): 293–296. doi: 10.1038/nsmb.2804.

Shapiro, J.A. 2022. Evolution: A View from the 21st Century. Cognition Press, Chicago.

Shen, G., W. Sun, Z. Chen, L. Shi, J. Hong, et al. 2022. Plant GDSL Esterases/Lipases: Evolutionary, Physiological and Molecular Functions in Plant Development. Plants 11(4): 468. doi: 10.3390/plants11040468.

Stewart, B. a., R. l. Baumhardt, and S. r. Evett. 2010. Major Advances of Soil and Water Conservation in the U.S. Southern Great Plains. Soil and Water Conservation Advances in the United States. John Wiley & Sons, Ltd. p. 103–129

Su, G., P. Madsen, M.S. Lund, D. Sorensen, I.R. Korsgaard, et al. 2006. Bayesian analysis of the linear reaction norm model with unknown covariates. J Anim Sci 84(7): 1651–1657. doi: 10.2527/jas.2005-517.

Su, Y., G.L.N. Ngea, K. Wang, Y. Lu, E.A. Godana, et al. 2024. Deciphering the mechanism of E3 ubiquitin ligases in plant responses to abiotic and biotic stresses and perspectives on PROTACs for crop resistance. Plant Biotechnology Journal 22(10): 2811–2843. doi: 10.1111/pbi.14407.

Sultan, S.E. 1987. Evolutionary Implications of Phenotypic Plasticity in Plants. In: Hecht, M.K., Wallace, B., and Prance, G.T., editors, Evolutionary Biology: Volume 21. Springer US, Boston, MA. p. 127–178

Tack, J., A. Barkley, and L.L. Nalley. 2015. Effect of warming temperatures on US wheat yields. Proceedings of the National Academy of Sciences 112(22): 6931–6936. doi: 10.1073/pnas.1415181112.

Therneau, T., and B. Atkinson. 1999. rpart: Recursive Partitioning and Regression Trees. : 4.1.23. doi: 10.32614/CRAN.package.rpart.

Thornton, P.E., M.M. Thornton, B.W. Mayer, N. Wilhelmi, Y. Wei, et al. 2014. Daymet: Daily Surface Weather Data on a 1-km Grid for North America, Version 2. Oak Ridge National Lab. (ORNL), Oak Ridge, TN (United States).

Turner, M.K., Y. Jin, M.N. Rouse, and J.A. Anderson. 2016. Stem Rust Resistance in ‘Jagger’ Winter Wheat. Crop Science 56(4): 1719–1725. doi: 10.2135/cropsci2015.11.0683.

Umate, P., R. Tuteja, and N. Tuteja. 2010. Genome-wide analysis of helicase gene family from rice and Arabidopsis: a comparison with yeast and human. Plant Mol Biol 73(4): 449–465. doi: 10.1007/s11103-010-9632-5.

USDA-NASS. 2024. United States Department of Agriculture - National Agricultural Statistics Service. https://quickstats.nass.usda.gov/ (accessed 16 August 2024).

Valladares, F., E. Gianoli, and J.M. Gómez. 2007. Ecological limits to plant phenotypic plasticity. New Phytologist 176(4): 749–763. doi: 10.1111/j.1469-8137.2007.02275.x.

Vierstra, R.D. 2012. The Expanding Universe of Ubiquitin and Ubiquitin-Like Modifiers. Plant Physiology 160(1): 2–14. doi: 10.1104/pp.112.200667.

Wang, X., X. Chen, Z. Liu, S. Tang, L. Zhang, et al. 2023. Genome-wide identification and functional characterization of pectin methylesterase inhibitors associated with male sterility in wheat. Environmental and Experimental Botany 212: 105383. doi: 10.1016/j.envexpbot.2023.105383.

Wang, X., X. Yao, A. Zhao, M. Yang, W. Zhao, et al. 2021. Phosphoinositide-specific phospholipase C gene involved in heat and drought tolerance in wheat (Triticum aestivum L.). Genes Genom 43(10): 1167–1177. doi: 10.1007/s13258-021-01123-x.

Wang, J., and Z. Zhang. 2021. GAPIT Version 3: Boosting Power and Accuracy for Genomic Association and Prediction. Genomics, Proteomics & Bioinformatics 19(4): 629–640. doi: 10.1016/j.gpb.2021.08.005.

Wang, H., S. Zou, Y. Li, F. Lin, and D. Tang. 2020. An ankyrin-repeat and WRKY-domain-containing immune receptor confers stripe rust resistance in wheat. Nat Commun 11(1): 1353. doi: 10.1038/s41467-020-15139-6.

Wei, J., W. Shao, X. Liu, L. He, C. Zhao, et al. 2022. Genome-wide identification and expression analysis of phospholipase D gene in leaves of sorghum in response to abiotic stresses. Physiol Mol Biol Plants 28(6): 1261–1276. doi: 10.1007/s12298-022-01200-9.

Wenig, U., S. Meyer, R. Stadler, S. Fischer, D. Werner, et al. 2013. Identification of MAIN, a factor involved in genome stability in the meristems of Arabidopsis thaliana. The Plant Journal 75(3): 469–483. doi: 10.1111/tpj.12215.

West-Eberhard, M.J. 2003a. Environmental Modifications. In: West-Eberhard, M.J., editor, Developmental Plasticity and Evolution. Oxford University Press. p. 794

West-Eberhard, M.J. 2003b. Material for a Synthesis. In: West-Eberhard, M.J., editor, Developmental Plasticity and Evolution. Oxford University Press. p. 794

Williamson, V.M., V. Thomas, H. Ferris, and J. Dubcovsky. 2013. An Aegilops ventricosa Translocation Confers Resistance Against Root-knot Nematodes to Common Wheat. Crop Science 53(4): 1412–1418. doi: 10.2135/cropsci2012.12.0681.

Woltereck. 1909. Weitere experimentelle Vntersuchungen iiber Artverandenmg, spcziell fiber das Wesen quantitativer Artunterschiede bei Daphniden. Verhandlungen der Deutschen Zoologischen Gesellschaft 19: 110–172.

Wu, Y., W. Wang, Q. Li, G. Zhang, X. Zhao, et al. 2020. The wheat E3 ligase TaPUB26 is a negative regulator in response to salt stress in transgenic *Brachypodium distachyon*. Plant Science 294: 110441. doi: 10.1016/j.plantsci.2020.110441.

Xue, S., J.A. Kolmer, S. Wang, and L. Yan. 2018. Mapping of Leaf Rust Resistance Genes and Molecular Characterization of the 2NS/2AS Translocation in the Wheat Cultivar Jagger. G3 Genes|Genomes|Genetics 8(6): 2059–2065. doi: 10.1534/g3.118.200058.

Yang, Y., A. Amo, D. Wei, Y. Chai, J. Zheng, et al. 2021. Large-scale integration of meta-QTL and genome-wide association study discovers the genomic regions and candidate genes for yield and yield-related traits in bread wheat. Theor Appl Genet 134(9): 3083–3109. doi: 10.1007/s00122-021-03881-4.

Yang, X., K. Wang, Y. Bu, F. Niu, L. Ge, et al. 2022. Genome-wide analysis of *GELP* gene family in wheat and validation of *TaGELP073* involved in anther and pollen development. Environmental and Experimental Botany 200: 104914. doi: 10.1016/j.envexpbot.2022.104914.

Zan, T., L. Li, J. Li, L. Zhang, and X. Li. 2020. Genome-wide identification and characterization of late embryogenesis abundant protein-encoding gene family in wheat: Evolution and expression profiles during development and stress. Gene 736: 144422. doi: 10.1016/j.gene.2020.144422.

Zhang, L.-Y., D.-C. Liu, X.-L. Guo, W.-L. Yang, J.-Z. Sun, et al. 2010. Genomic Distribution of Quantitative Trait Loci for Yield and Yield-related Traits in Common Wheat. Journal of Integrative Plant Biology 52(11): 996–1007. doi: 10.1111/j.1744-7909.2010.00967.x.

Zhang, N., Y. Yin, X. Liu, S. Tong, J. Xing, et al. 2017. The E3 Ligase TaSAP5 Alters Drought Stress Responses by Promoting the Degradation of DRIP Proteins. Plant Physiol. 175(4): 1878– 1892. doi: 10.1104/pp.17.01319.

Zhang, L., H. Zhao, N. Wan, G. Bai, M.B. Kirkham, et al. 2024. An unprecedented fall drought drives Dust Bowl–like losses associated with La Niña events in US wheat production. Science Advances 10(31): eado6864. doi: 10.1126/sciadv.ado6864.

Zhao, X., F. Goher, L. Chen, J. Song, and J. Zhao. 2023. Genome-Wide Identification, Phylogeny and Expression Analysis of Subtilisin (SBT) Gene Family under Wheat Biotic and Abiotic Stress. Plants 12(17): 3065. doi: 10.3390/plants12173065.

Zhao, C., B. Liu, S. Piao, X. Wang, D.B. Lobell, et al. 2017. Temperature increase reduces global yields of major crops in four independent estimates. Proceedings of the National Academy of Sciences 114(35): 9326–9331. doi: 10.1073/pnas.1701762114.

Zhao, X.-Y., J.-G. Wang, S.-J. Song, Q. Wang, H. Kang, et al. 2016. Precocious leaf senescence by functional loss of PROTEIN S-ACYL TRANSFERASE14 involves the NPR1-dependent salicylic acid signaling. Sci Rep 6(1): 20309. doi: 10.1038/srep20309.

Zhu, T., L. Wang, H. Rimbert, J.C. Rodriguez, K.R. Deal, et al. 2021. Optical maps refine the bread wheat Triticum aestivum cv. Chinese Spring genome assembly. The Plant Journal 107(1): 303–314. doi: 10.1111/tpj.15289.

